# WT and A53T *α*-synuclein systems: Melting Diagram and its new interpretation

**DOI:** 10.1101/724922

**Authors:** M. Bokor, Á. Tantos, P. Tompa, K.-H. Han, K. Tompa

## Abstract

Parkinson’s disease is connected with abnormal *α*-synuclein (*α*S) aggregation. Energetics of potential barriers governing motions of hydration water is examined. Information about the distributions and heights of potential barriers is gained by a thermodynamical approach. The ratios of the heterogeneous water-binding interfaces measure proteins’ structural disorder. All *α*S forms possess secondary structural elements though they are intrinsically disordered. Monomers are functional at the lowest potential barriers, where mobile hydration water exists, with monolayer coverage of mobile hydration. The *α*S monomer contains 33% secondary structure and is more compact than a random coil. A53T *α*S monomer has a more open structure than the wild type. Monomers realize all possible hydrogen bonds. Half of the mobile hydration water amount for monomers is missing in *α*S oligomers and *α*S amyloids. Oligomers are ordered by 66%. Mobile water molecules in the first hydration shell of amyloids are the weakest bound compared to other forms. Wild type and A53T amyloids show identical, low-level hydration, and are considered as disordered to 75%.

**Statement of Significance:** Aggregation of α-synuclein into oligomers, amyloid fibrils is a hallmark of Parkinson’s disease. A thermodynamic approach provides information on the heterogeneity of protein-water bonds in the wild type and A53T mutant monomers, oligomers, amyloids. This information can be related to ratios of heterogeneous water-binding interfaces, which measure the proteins’ structural disorder. Both α-synuclein monomers are intrinsically disordered. The monomers nevertheless have 33% secondary structure. They are functional as long as mobile water molecules surround them. They realize every possible H-bonds with water. Oligomers are like globular proteins with 66% ordered structure. Amyloids are disordered to 75% and are poorly hydrated with loosely bound water. Their hydration is identical. Oligomers, amyloids have only half as much hydrating mobile water as monomers.

## Introduction

WT and A53T mutant *α*S variants were investigated in the forms of monomers, oligomers, and amyloid fibrils. We compared previously details of the hydration of wild type *α*-synuclein (WT *α*S) and its A53T mutant by a combination of wide-line ^1^H NMR, differential scanning calorimetry, and molecular dynamics simulations (1). We interpret in this work, the wide-line ^1^H NMR results in the thermodynamic aspect, which was introduced in our earlier publications (2, 3). This point of view can be used to get information on the energetics of mobile-hydration-water binding properties of the proteins. The information obtained in this work includes the distributions of the potential barriers, which control the motion of the mobile hydration water. The term mobile hydration water refers to reorienting water molecules bound to the proteins also. Through the distribution of the potential barriers, information is gained about the strength of the protein-water bonds. The distribution of the different types of potential energies (different in magnitude and heterogeneity) shed light on the disordered or ordered global nature of the protein. The amounts of bound mobile water in different states were identified. So, quantities of water bound to heterogeneous or homogeneous parts of the solvent-accessible surface (SAS) of the proteins were distinguished and measured.

The studied proteins play an essential role in Parkinson’s disease (PD), which is an age-related neurodegenerative disorder diagnosed by tremor at rest, rigidity, and bradykinesia symptoms. It affects more than 5% of the aged population worldwide (above age 85) and is one of the biggest challenges of modern society (4). Aggregation of the *α*S protein is thought to be a critical step in the pathogenesis of PD and several other neurodegenerative disorders. Human *α*S is a 140-residue, highly conserved presynaptic protein that is abundant in various regions of the brain (5, 6). Structurally, purified *α*S is a monomeric, intrinsically disordered protein (IDP) at neutral pH (1, 7). The phrase IDP refers to proteins without fixed three-dimensional structures under physiological conditions. Due to their specific amino acid composition, IDPs, in general, have a unique combination of low overall hydrophobicity and large net charge (12). Though *α*S is an IDP, it shows secondary structure to a limited extent (8–10) and it is slightly more compact than a random coil (9, 11). Furthermore high-resolution NMR analysis revealed that it exhibits a region with a preference for helical conformation (11).

Disease-related mutant forms of *α*S have been identified earlier. Alanine-to-threonine substitution at the 53th amino acid residue entail inverse preferences of alanine to form helices and of threonine to support *α*-sheet structures, which are crucial for amyloid fibril formation. Accordingly, the A53T variant, which is an early onset mutant, was predicted to be more likely to form β-sheet structure than WT *α*S (13, 14). It forms fibrils the fastest in vitro (13), and WT is the slowest in fibril formation of all the WT and familial mutant variants (A30P, E46K, A53T). *α*S oligomers, the precursors of fibrils is more toxic than mature fibrils. A53T mutation has been shown to alter the neurotoxicity and aggregation properties of the WT *α*S protein (14). Dynamics of the proteins play an important role in the process of amyloid fibril formation (15). The WT and A53T *α*S monomers exhibit almost identical structural properties and conformational behaviour (1,16,17). The oligomers exist in a range of sizes, with different extents and nature of β-sheet content and exposed hydrophobicity (18). They contain 35±5% of β-sheet structure, as opposed to 0% in the soluble *α*S monomers and 65±10% in the *α*S fibrils (19). We have shown earlier that the oligomeric form is ordered similarly to the globular proteins (1). The molecular mechanisms underlying *α*S aggregation remain unknown (14, 20).

Wide-line ^1^H NMR, differential scanning calorimetry, and molecular dynamics simulations suggest a hydrate shell of *α*S compatible with largely disordered states (1). The A53T mutant displays a somewhat higher level of hydration than WT *α*S, suggesting a bias to more open structures, favourable for protein-protein interactions leading to amyloid formation. The differences between WT and A53T *α*S disappear in the amyloid state, suggesting the same surface topology, irrespective of the initial monomeric state. Molecular dynamics simulations showed that the wide-line ^1^H NMR results describe the first hydration layer (1). We have demonstrated that wide-line ^1^H NMR spectrometry can distinguish point-mutants based on the binding heterogeneity of the protein surface (2, 3). The NMR measurements presented in reference (1), were not focused on the structures of the proteins. They presented an interpretation process based on the measurement of the mobility of hydration water. The interpretation of the resulting melting diagrams (MD, amount of mobile hydration vs. normalized fundamental temperature) informs about the thermodynamic nature of the SAS of the protein. We present here an energetic description of the potential barriers governing motions of water molecules in the hydration shell of *α*S (2, 3). Wide energy distributions of the potential barriers are experienced, and the gained information is not just an average value. The potential barriers reflect the protein-water interactions and vary in chemical and topological properties of the interactive SAS of the protein. The measurement and the interpretation process we applied for *α*S monomer variants make also possible to investigate molecular interactions. Oligomerization and amyloid formation play a role in disease occurrence. The degree of hydrophilicity of the protein’s interaction sites determines quantity of hydration. We drew quantitative conclusions about the ratios of the ordered and disordered, more solvent-exposed surface regions of proteins and the extent of the latter, as well as the energy relations of the protein-water bonds (3). Hydrogen bonds between protein and water play a dominant role in the formation of the first hydrate shell.

We introduced normalized fundamental temperature, and an energy scale was set up in the form of normalized fundamental temperature being scaled with the molar enthalpy of fusion for water. The normalization is made by the melting point of bulk water. The dynamic order parameter of the *MD, i.e.* the heterogeneity ratio characterizes protein-water-bond energy distribution. The usage of fundamental temperature helps to describe *MD* in the form of power series. The derivative form of the latter defines the number of water molecules that begin to move at a given potential barrier.

Our investigation is aimed at the exploration of the structural differences between WT and A53T *α*S variants, and furthermore at the investigation of the potential barrier distributions around the different *α*S variants at the actual thermal energy. Wide line ^1^H-NMR was used as a tool to record the *MDs* of the frozen protein solutions (2,21–23). The amount of mobile hydration water is calculated from the amplitude of the wide line ^1^H NMR signal. In this work, only a very short description is provided, for more details of the measuring method, see reference (3). Now, we can set up energy distributions of the potential barriers affecting the motion of hydration water, for better understanding and deeper insight.

## Materials and Methods

### Proteins

Expression and purification of recombinant human wild-type and A53T mutant *α*-synuclein in a pRK-172-based expression system was performed as described in (24). In sample preparation, the mass of lyophilized protein (without any further refinement) was measured and an appropriate amount of water was added to obtain the requested nominal concentrations, 50 mg/ml and 25 mg/ml. Ultra-pure double-distilled water was used as a solvent. Measurements were carried out on three identical samples prepared independently. Amyloid was prepared by preincubating protein samples at 3 °C for 24 h. The formation of amyloid was confirmed by measuring the fluorescence of 5 mM thioflavin T (ThT) added to an aliquot of the solution. All results where a mobile fraction of water, n(Tfn) is involved, are calculated for 50 mg/ml protein concentration.

### Wide-line NMR measurements

^1^H-NMR measurements and data acquisition were accomplished by a Bruker AVANCE III NMR pulse spectrometer at 82.6 MHz frequency, with a stability better than ±10^-6^. The inhomogeneity of the magnetic field was 2 ppm. The data points in the figures are based on spectra recorded by averaging the signals to reach a signal/noise ratio of 50. The extrapolation to zero time was done by fitting a stretched exponential. The principle and details of wide-line ^1^H NMR spectroscopy are given in refs. (22, 23).

Free induction decay signals (FIDs) were measured between −70 °C and +25 °C, following thermal equilibrium. The temperature was controlled by an open-cycle Janis cryostat with stability better than ±1 K. The phases of ice protons, protein protons, and mobile (water) protons are separated in the FID signal by large differences in their spin-spin relaxation rate. The portion of mobile proton (water) fraction is directly measured by the FID signal and Carr-Purcell-Meiboom-Gill (CPMG)-echo trains (in the temperature range where echoes could be detected at all) (22, 23).

Melting is understood in the frozen protein solutions here, as the appearance of a motionally narrowed component in the ^1^H NMR-spectrum, attributed to mobile water molecules (2). The melting diagram or curve gives the amount of meltwater - in this sense - as a function of temperature. The amount of meltwater, *n* is measured by the FID amplitude directly as a fraction of the total water content of the protein solution. It is converted into hydration, *h* as gram water per gram protein (Figs. 1–3). Hydration is an intensive quantity, independent of the size or concentration of the protein.

**Figure 1.**
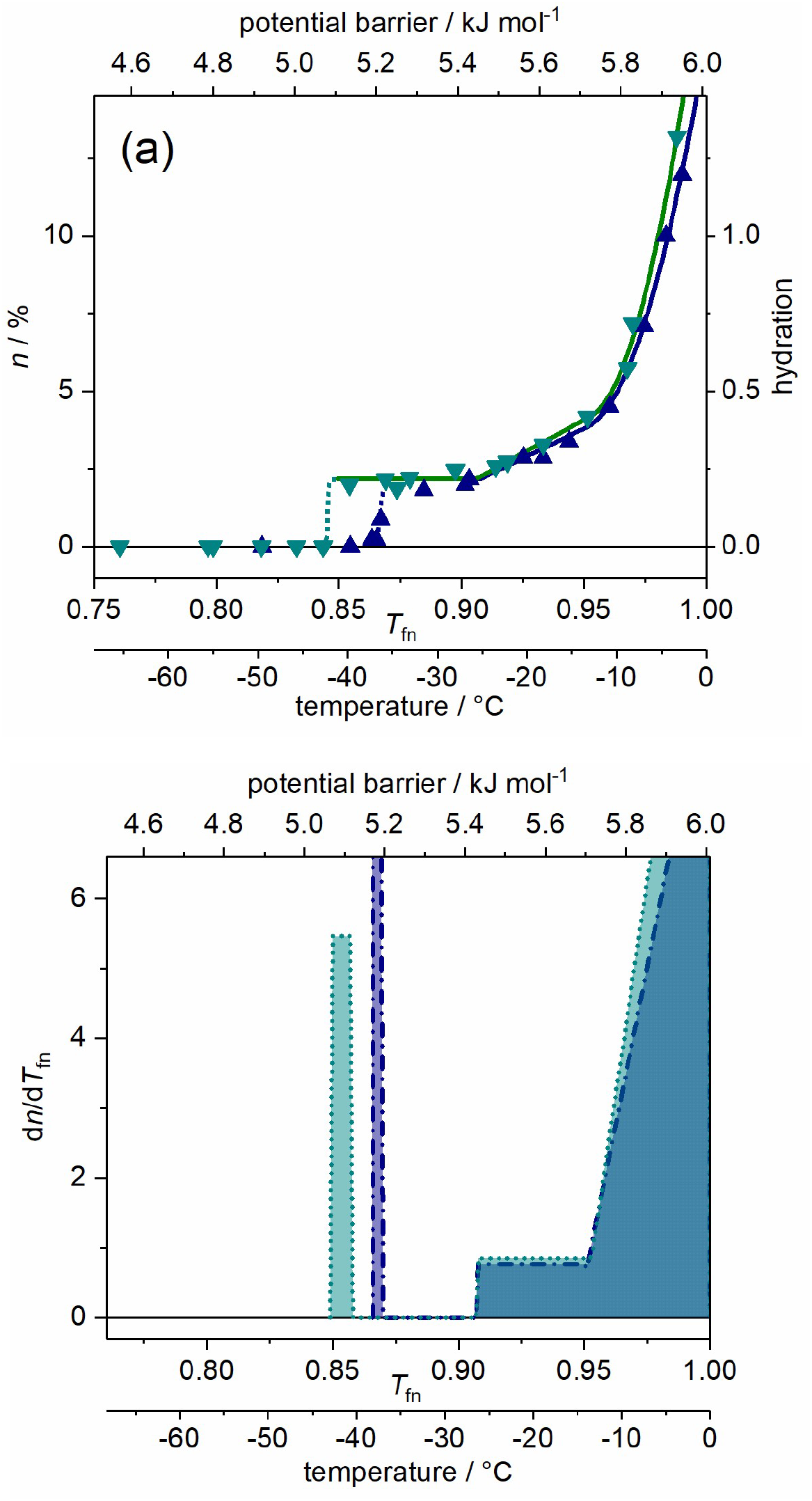
*α*S monomer, wild type (dark blue) and A53T mutant (cyan) variants dissolved in pure water. (*a*) Melting diagram. (*b*) Derivative of the melting diagram, *i.e.* potential barrier distribution related to moving hydration water. There are no reliable measured data in the range ^-^1-0 °C (0.995-1.00 *T*_fn_). Data are given for 50 mg ml^-1^ protein concentration.

**Figure 2.**
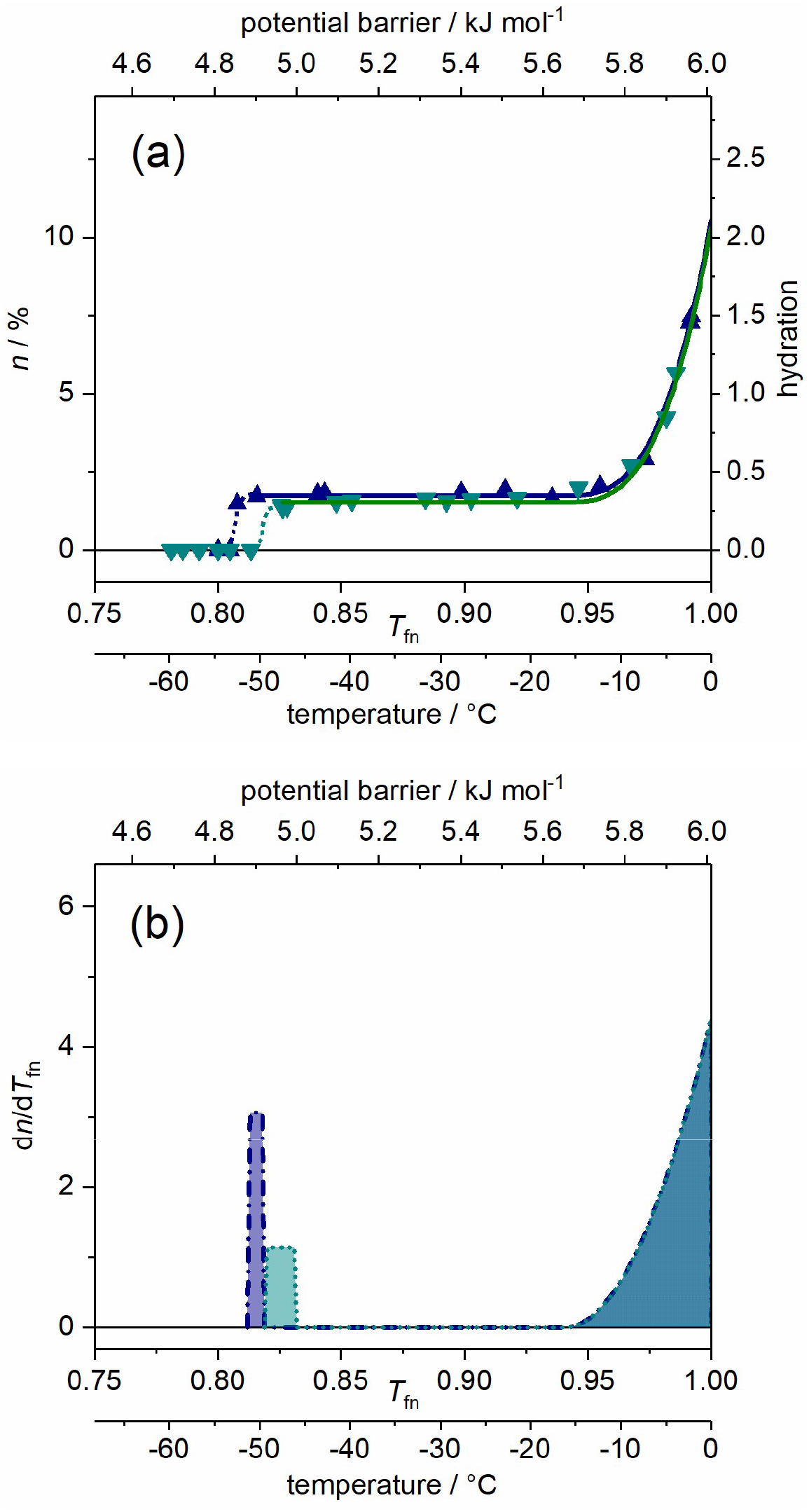
*α*S oligomer, wild type (dark blue) and A53T mutant (cyan) variants dissolved in pure water. (*a*) Melting diagram. (*b*) Derivative of the melting diagram, *i.e.* potential barrier distribution related to moving hydration water. There are no reliable measured data in the range ^-^1-0 °C (0.995-1.00 *T*_fn_). Data are given for 50 mg ml^-1^ protein concentration.

**Figure 3.**
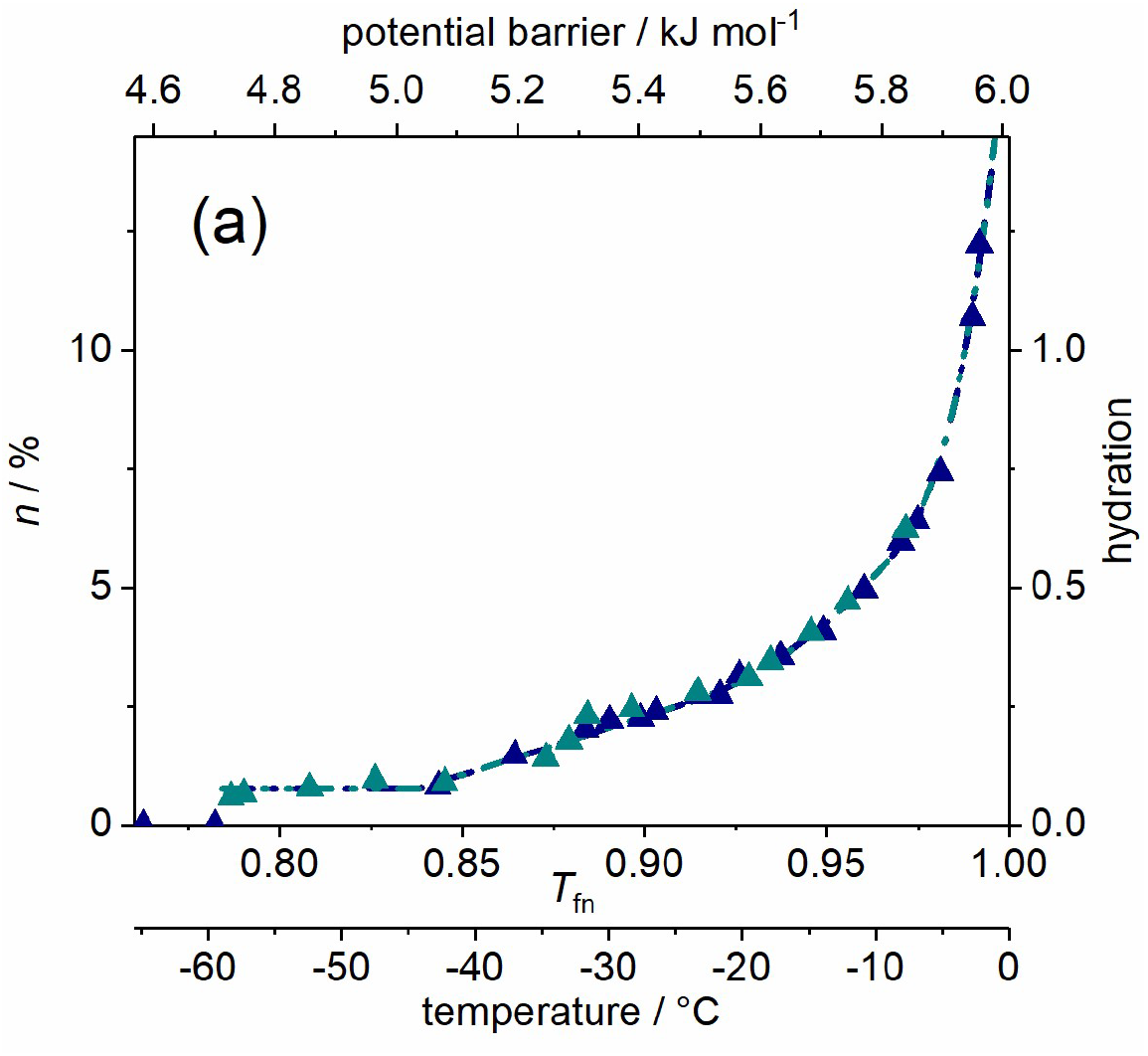
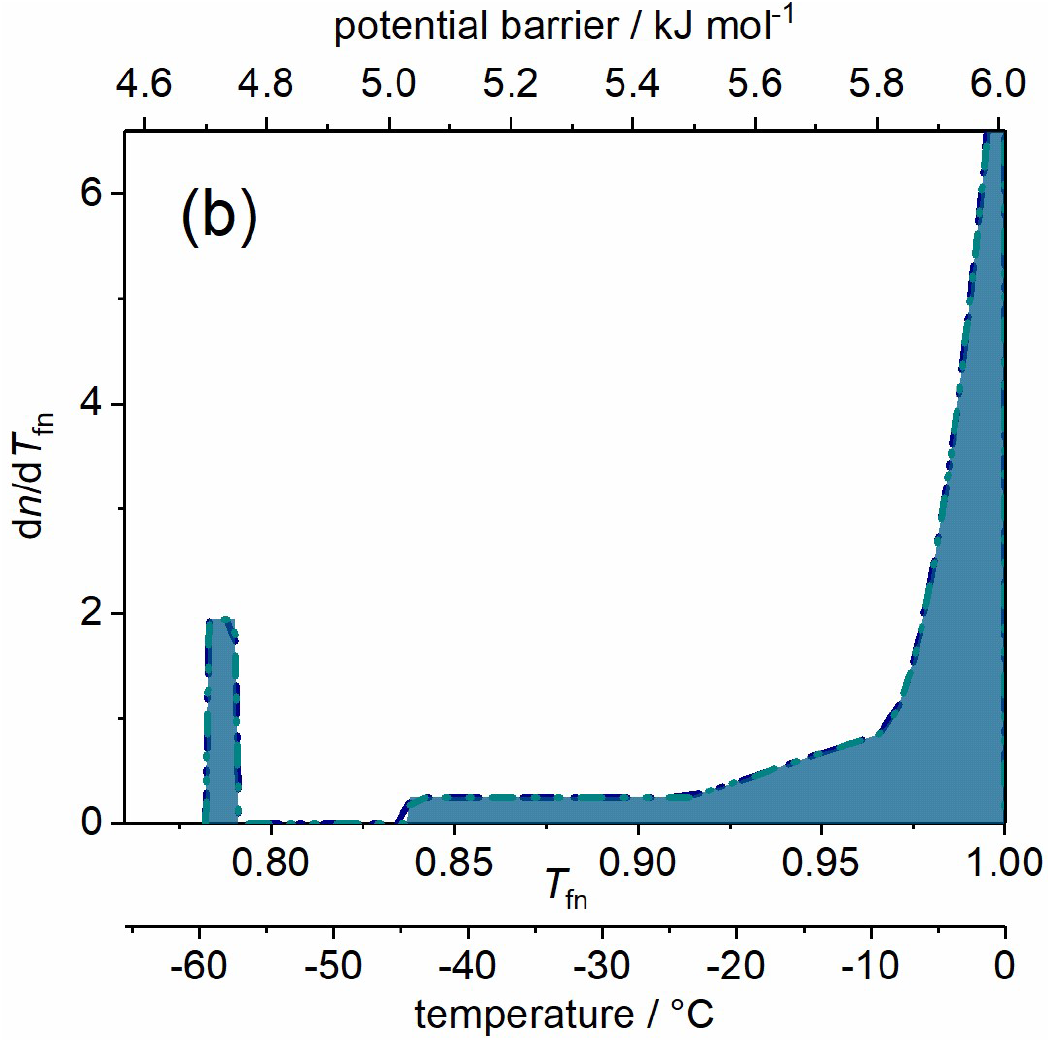
*α*S amyloid, wild type (dark blue) and A53T mutant (cyan) variants dissolved in pure water. (*a*) Melting diagram, the data are coincident for the two variants. (*b*) Derivative of the melting diagram, *i.e.* potential barrier distribution related to moving hydration water. There are no reliable measured data in the range ^-^1-0 °C (0.995-1.00 T_fn_). Data are given for 50 mg ml^-1^ protein concentration.

### Interpretation methods of the melting diagram

As temperature measurement, fundamental temperature is used, *T*_f_ = *k*_B_ *T* or *T*_f_ = *RT*, where *T* is absolute temperature. A normalized version of the fundamental temperature scale was introduced, normalized to the melting point of bulk ice, *T*_fn_ = *T*/273.15 K. The start of a molecular motion can be described in solids by the necessary thermal input as *V*_0_ = *constant·T* (25). This idea was applied to frozen aqueous protein solutions and was corrected dimensionally (2, 3). The *T*_fn_ scale is converted into an energy scale, scaled with water’s heat of fusion, 6.01 kJ mol^-1^ (26). The energy scale is given by the relation *E*_a_ = *cRT* on a molecular level, where *c* is a constant. No modelling assumptions are used in the interpretation. Molecular motion is considered to be rotation and not vibration.

Proteins can be characterized and categorized by the heterogeneity of the energy distribution of water binding, described by the heterogeneity ratio

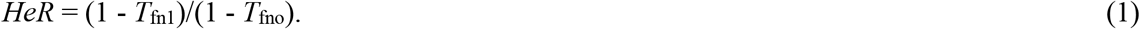

*T*_fno_ and *T*_fn1_ are start and endpoints of the temperature-independent section of the *MDs*. *HeR* provides the ratio of the heterogeneous binding interface, and it is a measure of the structural disorder of proteins. The *MDs* can be described analytically by a power series as

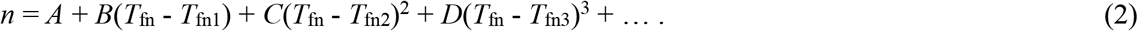

The derivative form gives the change of *n* and *DMD* (a derivative form of *MD*) is then

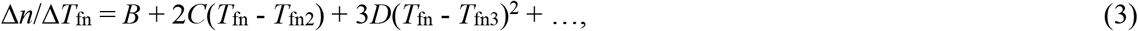

and highlights the peculiarities of the hydration melting process.

The ratio of heterogeneous protein-water bonds, *HeR*_n_ (3) is calculated with *n*_ho_, the number of water molecules in the first hydration layer and with the total number of water molecules in the whole of the heterogeneous region, *n*_he_ as

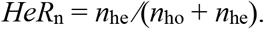

The measure of heterogeneity, *HeM* describes the heterogeneity of the protein-water bonds near *E*_a_ = 6.01 kJ mol^-1^ and gives fingerprint of each protein. The formula (2, 27) is related to the tangent function and to *DMD*,

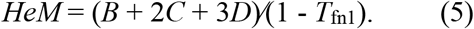

For more details on the evaluation and interpretation of ^1^H NMR data, see (2, 3) and the *Supporting Information* to (28).

## Results and Discussion

The first hydration layer on the protein surface is of particular importance for biological activity (29). A protein with its first hydration layer forms the biologically active entity (30). Hydration water moves on the picosecond time scale while NMR-processes are slower by an order of magnitude. The phenomenon of motional narrowing in NMR is a good indicator of mobile water.

Proteins do not interact directly with the bulk water phase. Their interactions are realized with a few bound (31) or non-freezing water layers. This fact rationalizes studying the properties of bound water. Bound or non-freezing water is referred here as mobile hydration water. Hydration water must be regarded not as a rigid shell of water molecules around the protein molecules, but as a fluctuating cloud of water molecules that interacts more or less strongly (even unfavourably) with the protein surface. The protein-water interaction energies are considered to equal with the relevant potential barrier values for the mobile hydration water. Wide energy distributions are experienced, and the gained information is not just an average value. The extent of the exposed hydrophobicity determines how strongly water is bound in the hydration shell of a protein (32). It is inversely proportional to bond strength. The motion of the hydration water molecules, which are the most loosely bound to the proteins is hindered by potential barriers very different in magnitude for the different forms of *α*S. The disordered nature of IDPs originates from a large content of hydrophilic amino acid residues compared to hydrophobic ones, preventing the hydrophobic collapse (33). If the amino acid composition of a protein is known, predictions can be made of the amount of bound water (31, 34).

### *α*-Synuclein Monomers

The WT and A53T variants of *α*S monomers dissolved in pure water show *MDs* characteristic of IDPs (Fig. 1) (1–3,21–23) as follows. (i) Mobile hydration water molecules appear around this proteins at a relatively high potential barrier value as the lowest (*E*_a,o_, Table 1). The actual Ea,o counts as especially high for *α*S monomers when compared to oligomer and amyloid *α*S forms or to globular proteins (2). *(ii)* Moreover, the *MDs* of IDPs have a steeply increasing trend with rising potential barrier values from *E*_a,1_ to *E*_a_ = 6.01 kJ mol^-1^ (0 °C; Fig. 1) (1–3,21–23).

**Table 1.** Parameter values for the polynomial relation (Eq. 1) describing mobile water fraction, *n*. The error in the last digit is given in parentheses.

| polymerization | monomer |  | oligomer |  | amyloid |
| --- | --- | --- | --- | --- | --- |
| αS variant | WT<sup>(a)</sup> | A53T | WT | A53T | WT, A53T |
| $A = n(T_{\text{fno}})^{(b)} = n(T_{\text{fn1}})$ | 0.22(4) | 0.22(4) | 0.0180(4) | 0.0157(4) | 0.0039(3) |
| $h(E_{a,o})^{(c)} = h(E_{a,1})$ | 0.44 (8) | 0.44(8) | 0.360(9) | 0.315(8) | 0.077(6) |
| $B$ | 0.38(5) | 0.43(3) | 0 | 0 | 0.12(1) |
| $C$ | 45(7) | $0.6(1) \cdot 10^2$ | 0 | 0 | 2.9(4) |
| $D$ | 0 | 0 | $4(1) \cdot 10^2$ | $4.1(7) \cdot 10^2$ | $10(2) \cdot 10^2$ |
| $T_{\text{fno}} / \text{kJ mol}^{-1}$ | 0.8662(9) | 0.854(5) | 0.808(3) | 0.820(6) | 0.784(2) |
| $E_{a,o} / \text{kJ mol}^{-1}$ | 5.206(6) | 5.13(3) | 4.85(2) | 4.93(4) | 4.71(1) |
| $t_o / ^\circ\text{C}^{(d)}$ | -36.5(2) | -40(1) | -53.5(8) | -49(2) | -58.9(6) |
| $T_{\text{fn1}}$ | 0.908(4) | 0.906(2) | 0.941(5) | 0.939(3) | 0.966(2) |
| $E_{a,1} / \text{kJ mol}^{-1}$ | 5.46(2) | 5.44(1) | 5.66(3) | 5.64(2) | 5.81(1) |
| $t_1 / ^\circ\text{C}$ | -25.2(1) | -25.7(5) | -16(1) | -16.6(9) | -9.2(6) |
| $T_{\text{fn2}}$ | 0.951(3) | 0.953(4) | — | — | 0.914(3) |
| $E_{a,2} / \text{kJ mol}^{-1}$ | 5.72(1) | 5.73(2) | — | — | 5.49(2) |
| $t_2 / ^\circ\text{C}$ | -13.3(8) | -13(1) | — | — | -23.5(9) |
| $T_{\text{fn3}}$ | — | — | 0.941(5) | 0.939(3) | 0.966(2) |
| $E_{a,3} / \text{kJ mol}^{-1}$ | — | — | 5.66(3) | 5.64(2) | 5.81(1) |
| $t_3 / ^\circ\text{C}$ | — | — | -16(1) | -16.9(9) | -9.2(6) |
| $n(T_{\text{fn2}})$ | 0.039(3) | 0.042(8) | — | — | 0.013(2) |
| $h(E_{a,2})$ | 0.77(6) | 0.77(3) | — | — | 0.26(4) |
| $n(T_{\text{fn3}})$ | — | — | 0.0174(6) | 0.0153(4) | 0.03(2) |
| $h(E_{a,3})$ | — | — | 0.35(1) | 0.306(8) | 0.5(4) |
| $n(T_{\text{fn}} = 1)$ | 0.164(5) | 0.20(6) | 0.11(1) | 0.11(3) | 0.09(3) |
| $h(E_a = 6.01 \text{ kJ mol}^{-1})$ | 3.3(1) | 4(1) | 2.1(2) | 2.1(6) | 1.8(5) |

<sup>(a)</sup> Wild type. <sup>(b)</sup> Amount of mobile water fraction at $T_{\text{fno}}$ . <sup>(c)</sup> Hydration at $E_{a,o}$ . <sup>(d)</sup> Temperature in units of degrees Celsius.

The *MDs* of WT and A53T *α*S monomers show rapidly changing trends as a function of potential barriers governing the motion of hydration water, except for a short constant region at the start of mobility (Fig. 1). The amount of mobile water molecules, *h* = 0.44(8) is enough to have functional proteins with approximately monolayer coverage at the smallest potential barriers below Ea,1, where constant *h* region can be found. The mobile water concentration given as hydration provides a macroscopic measure. This level of mobile hydration water equals to 3.5(6)·10^2^ H_2_O/protein on the microscopic scale. Hydration, *h* is the microscopic mobile water fraction, *n* measured by NMR multiplied by the mass of water and divided by the mass of protein in the solution, *h* = *n*·*m*_water_/*m*_protein_. It is a measure of concentration for aqueous protein solutions on macroscopic scale. Molecular dynamics simulations show that the measured behaviour corresponds to the first hydration layer of the structural ensemble of the protein (1). Earlier NMR measurements and molecular dynamics simulations for individual amino acid’s hydration yield 4(1)·10^2^ water molecules per amino acid sequence of the whole protein (34–36). The comparison between our results and the predicted values suggests that the solvent accessible amino acid residues are fully hydrated with mobile water molecules in the monomers as soon as water molecules are no longer rigid in ice phase. It should be noted that NMR sees rotational motions only and no translational motion is detected here. It can be concluded that the amount of homogeneously, the weakest bound hydration water, *n*(*T*_fno_) = *n*(*E*_a,o_) is as high as if every possible hydrogen bond site were occupied by a water molecule. This refers to open structure.

Water in interaction with A53T *α*S gets in motion at a lower potential barrier compared to the WT variant (Fig. 1). This fact indicates that A53T *α*S interact with water weaker, its protein-water bonds are looser. The constant-value *h* regions at low potential barriers (from *E*_a,o_ to *E*_a,1_, Table 1) indicate the presence of secondary structures even though these proteins are IDPs. The occurrence of secondary structures in IDPs is not unexpected (37). These regions appear as spikes and subsequent zero values in DMDs. Consequently, only one type of potential barrier or protein-water interaction is responsible for the motion of water molecules in this energy region. The type of the interaction is not van der Waals here, because it lacks continuous distance dependence; this behaviour suggests the formation of hydrogen bonds.

The plateau region (constant value of *h*) is wider for A53T *α*S but the *E*_a,1_ values are the same for both variants (Table 1). This means that A53T *α*S has a wider spread of non-occupied protein-water interaction energy levels and the potential barrier gap is wider. The explanation for this can be that the A53T mutation results in a slight enhancement of the region around mutation site with a preference to form extended conformations (38).

WT and A53T *α*S monomer proteins are intrinsically disordered and that is why their *MDs* show rapidly changing thermal trends above *E*_a,1_ (Fig. 1). Solvent-exposed amino acid residues result in a solvent accessible surface of the protein highly heterogenic by energy. Elevation of *MD* for *α*S monomers in the linear section (*E*_a,1_ < *E*_a_ < *E*_a,2_) is more than three times steeper than in the case of *α*S amyloid (parameter B, Table 1). The introduction of heterogeneous distribution of potential barriers at *E*_a,1_ is connected to general change in the motional state of the hydration water. As a blanket result, it was found for both globular proteins and IDPs that dynamic contributions to the quasi-elastic neutron scattering spectra (30) all change their thermal trend around *E*_a,1_. At the higher hydration levels, not only hydrogen bonds are responsible for water-protein interactions but van der Waals forces also are acting. A section of quadratic trend was detected in the *MDs* for both *α*S monomer variants above *E*_a,2_ (Fig. 1).

Fibril formation is inhibited by intramolecular interactions in the WT *α*S structure and these interactions are greatly destabilized by the A53T mutation (39), shifting the conformational ensemble of *α*S to a more open state (40). In accordance with this statement, wide line ^1^H NMR detected that A53T *α*S has a higher level of hydration than the WT variant (Fig. 1a) meaning that the mutation resulted in an overall more open structure. The *α*S monomers have the highest ratio of exposed hydrophilic groups with five times more moving water in their first hydration shell than the *α*S amyloids.

The *DMD* (Fig. 1*b*) provides a quantitative measure of the energetic heterogeneity of the potential barriers affecting the mobility of hydration water molecules. The *α*S monomer-water interaction energy is most diversified. This can be understood as resulting from the open structure of the monomers, with many different types of hydrophilic groups in their water accessible surface. The difference between the *MD* of the WT and A53T *α*S monomers is very minute (Fig. 1), though the A53T mutant is steadily more heterogeneous with higher hydration levels.

WT and A53T *α*S monomers reach the far highest level of hydration compared to oligomers and amyloids at the melting point of bulk water, *h*(*E*_a_ = 6.01 kJ mol^-1^, monomer) = 3.3(7), substantiating the most open structure for them. The observed *HeR* (Table 2) is not 1 as expected if *α*S monomers were totally disordered but it scores 33(4)% homogeneous potential barrier distribution for WT and A53T *α*S. This fact is in accordance with the structural ensemble of *α*S being more compact than it is expected for a random coil state (41, 42). The ratios of heterogeneously bound mobile water fractions to the sum of both heterogeneously and homogeneously bound water, *HeR*_n_*s* are equal within experimental error for all of the *α*S variants, except *α*S amyloids (Table 2). *α*S monomers have, nevertheless, little higher *HeR*_n_ values than *α*S oligomers and *α*S amyloids are characterized by far the highest ratio of heterogeneously bound mobile hydration water. *HeR*_n_ = 0.87(1) is calculated as average for the monomer and the oligomer variants, indicating high heterogeneity in the nature of the bonds formed by these *α*S variants. By this ratio, *α*S amyloids show the most heterogeneously bound water molecules in this region near to the melting point of bulk water. The amount of heterogeneously bound water for the monomers, *h*_he_^monomer^ = 2.8(2) is much higher than for oligomers (*h*_he_^oligomer^ = 1.760(4)) and amyloids (*h*_he_^amyloid^ = 1.68(9)). This relation of hhe values reflects that **α**S monomers are IDPs, the *α*S oligomers and *α*S amyloids are significantly more structured. *h*_he_ serves as nominator in the expression of *HeR*_n_. The measure of heterogeneity, *HeM* characterizes the extent of heterogeneity of the protein-water interactions close to *E*_a_ = 6.01 kJ mol^-1^. WT and A53T *α*S monomers show *HeM* values an order of magnitude smaller than *α*S oligomers and *α*S amyloids. The hydration of WT and A53T *α*S monomers is already heterogenic even at *E*_a,2_ and it doesn’t elevate intensively when approaching *E*_a_ = 6.01 kJ mol 1.

**Table 2.**
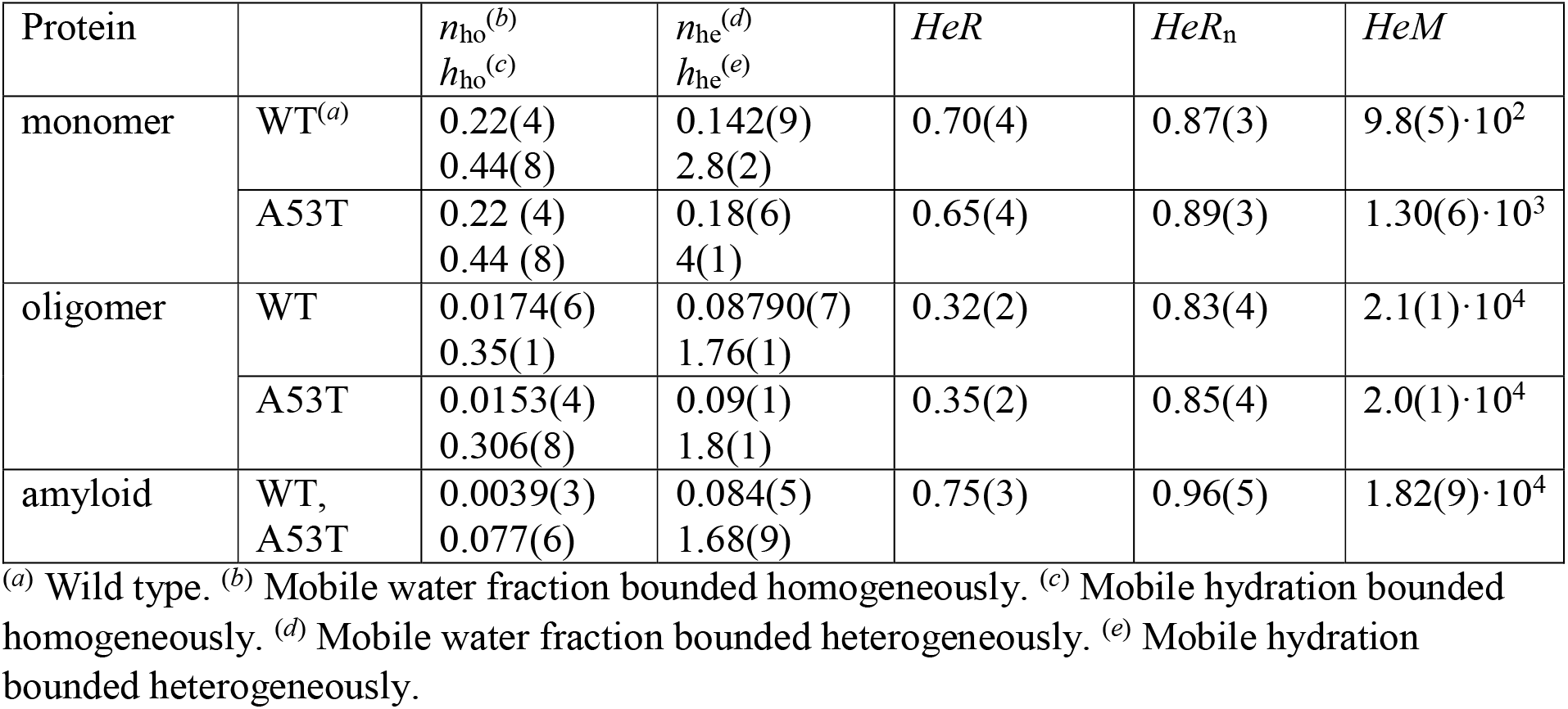
Homogeneously or heterogeneously bound amounts of water, and dynamic parameters. The error in the last digit is given in parentheses.

### α-Synuclein Oligomers

Features common to the different types of *α*S oligomers include β-sheet structures, high content of solvent-exposed hydrophobic regions and globular or tubular morphology (18, 43). The general surface of the *α*S oligomers is essentially the same irrespective of their size, which varies. This uniformity is assumed because there can be seen but only one type of oligomers below *E*_a,3_, according to the *MDs*.

The oligomer forms of WT and A53T *α*S both show *MDs* similar to those of globular proteins (2). This manifests itself in attributes as (*i*) the *E*_a,o_ is much lower than in the case of the IDP monomers; (*ii*) the wide temperature-independent region in their *MDs*; and (*iii*) steep rise in the hydration level shows up at high potential barriers near to *E*_a_ = 6.01 kJ mol^-1^.

The *MDs* of the oligomeric *α*S forms (Fig. 2a) are dominated by a wide constant hydration region. This is a general characteristic of ordered structure present in the *α*S oligomers. The very wide constant *h* region can be caused by the greater extent of solvent exposed hydrophobic surface (18). There is only one constant *MD* region with one single *h* value. This fact means that there are no water molecules, the motion of which is controlled by potential barriers with energies *E*_a,1_ < *E*_a_ < *E*_a,3_. The constant *h* region in the *MD* of WT *α*S oligomers is wider than that in the *MD* of A53T *α*S oligomers. Both are far wider than that of the IDP *α*S monomer forms. One single potential barrier (or a very narrow distribution of it) is characteristic for the motion of the *h*(*E*_a,1_) amount of mobile water, meaning a homogeneous energy distribution of potential barriers for them. These water molecules are homogeneous in their bonding properties since no others start to move in a very broad potential energy range.

The *α*S oligomers have a significant surface ratio of hydrophilic nature, which manifests itself in the wide heterogeneous *MD* section just below *E*_a_ = 6.01 kJ mol^-1^ (the growth can be described with a cubic term in the *MDs*). Our results may reflect the variety in oligomer *α*S size as the potential barrier range with an intensively growing amount of mobile water is relatively wide compared to globular proteins. That is, a very heterogeneous distribution of the actual potential barriers can be found for the water molecules starting to move in this energy range. Not only the amount of mobile hydration water grows rapidly, while it approaches the melting point of bulk water, but the intensity of the growth also increases.

The small *HeR* values of the WT and A53T *α*S oligomers mean that they are remarkably more ordered than the WT and A53T *α*S monomers (Table 2). The mobility in their hydration shell is more limited than for the monomers. *α*S oligomers show homogeneous protein-water interactions till *E*_a,1_ = *E*_a,3_ and their hydration is low at this potential barrier level, *h*(*E*_a,3_) = 0.32(2) (on the average). WT *α*S oligomers have higher *HeM* values than A53T *α*S oligomers and *α*S amyloids, which means that their bonding network is more heterogeneous close to the melting point of bulk water. The latter two have very similar *HeM* values. In a narrow energy spread of Δ*E*_a_ = 0.363(5) kJ mol^-1^, *α*S oligomers reach a relatively high hydration level by *E*_a_ = 6.01 kJ mol^-1^, *h*(*E*_a_ = 6.01 kJ mol^-1^) = 2.1(4). This behaviour also can be a sign of their variety in size. The growth of hydration is Δ*h* = 1.760(4) for *α*S oligomers in the region of heterogeneous elevation, at *E*_a,3_ < *E*_a_ < 6.01 kJ mol^-1^, that equals the size of the heterogeneously bound water fraction, *n*_he_ (Table 2). *HeM* values of oligomers are the highest indicating an intensive growth of bond heterogeneity in this region, starting from a low level. Hydrations of *α*S oligomers at *E*_a_ = 6.01 kJ mol^-1^ are much lower than that of *α*S monomers here and are a little greater than that of *α*S amyloids.

### α-Synuclein Amyloids

Fibrillary conformation of *α*S is largely composed of parallel β-sheet structures (18, 44). The weak bonds holding the parallel β-sheets together cause that the water molecules in the hydrate shell of the *α*S amyloids can be mobilized the most easily. Water bound to *α*S oligomer molecules requires higher energy to move, and the hydration water of the *α*S monomers can be moved at the highest energy cost. The mutations associated with familial PD (e.g. A53T) increase the propensity of *α*S monomer to form insoluble aggregates and produce morphologically distinct aggregate species according to literature data (45–49). The amyloid state shares distinctive structural features that are largely independent of the protein sequence (50, 51). We could see identical *MDs* of the WT and A53T amyloid variants (Fig. 3) in accordance with this feature and therefore, they are discussed unified.

The quantity of mobile hydration water at the start of motion is by far the smallest in amyloids compared to other *α*S states so it can be assumed that they have the greatest extent of exposed hydrophobicity. They have *h*(*E*_a,o_) = 0.077(6), a value that is approximately six times smaller than for *α*S monomers and more than four times smaller than for *α*S oligomers (Table 1, Figs. 1–3). The quantity of mobile hydration reaches the level needed for functionality at *E*_a,2_ only (minimally *h* = 0.2 is needed for protein functioning (52, 53)). The very low Ea,o is a characteristic feature of amyloids, even the globular proteins show higher potential barrier values as the lowest where mobile hydration water exists (2). *α*S amyloids have long cylindrical structures with water-filled interfaces (54–59). This implies that significant solvent space exists within the fibrils, which is attributed to the *α*S molecules within the fibrils having a distribution of conformations. The water inside the structure cannot be seen as mobile by wide line ^1^H NMR due to limited mobility. This fact can be responsible for the small amounts of mobile hydration water around the *α*S amyloids.

The low *E*_a,o_ and the subsequent zero value region in the *DMD* refer to highly hydrophobic patches of the solvent accessible surface of the *α*S amyloid fibrils. The constant mobile hydration region in the *MD* is half as wide as in the *α*S oligomers, but it is still twice as wide as in the *α*S monomers. The hydrophobic patches relate to secondary structures. Mature fibrils contain several protofilaments with a cross β-structure, in which individual β-strands run perpendicular to the fibril axis (60–64). Only specific segments of the chain are incorporated into the cross-β structure, with the remaining of the chain being external to the core elements of the structure. The central portion of *α*S corresponding to amino acid residues 31-109 (half the size of the holoprotein) constitutes the core of the *α*S amyloid filaments (19).

The abundant solvent-exposed side chains (15) are most possibly responsible for the mainly heterogeneous potential barrier distribution affecting the motion of the mobile hydration water at *E*_a,1_ < *E*_a_ < 6.01 kJ mol^-1^ (Fig. 3), and therefore they are responsible for the heterogeneous protein-water interactions. Further properties that increase heterogeneity are the large amplitudes of the side chain motions (larger in the fibril state than in the monomeric state, the amyloid state is entropically favourable (15)) and heterogeneity is also increased by the fibrils having a distribution of conformations. The *α*S amyloid form can be considered as disordered to 75(3)% or conversely, it is ordered to a measure of 25(3)% regarding the nature of its solvent accessible surface (*HeR* value, Table 2). *HeM* values for *α*S amyloids are of very similar value to that for oligomers and are much higher than monomers’ *HeM.* Bonding heterogeneity and *h*(*E*_a_ = 6.01 kJ mol^-1^) levels are quite similar for *α*S amyloid and oligomer variants. The main difference between these two aggregated species is that *α*S oligomers are ordered in contrast with disordered *α*S amyloids.

The hydration value at the melting point of bulk water, *h*(*E*_a_ = 6.01 kJ mol^-1^) for the *α*S amyloids is comparable to that of the *α*S oligomeric forms, but it is significantly lower than in the case of *α*S monomers (Table 1). This relation of the relative levels means that not only the *α*S oligomers, but also the *α*S amyloids have a compact structure compared to *α*S monomers. In an assembled *α*S amyloid, long N-terminal and C-terminal segments remain unprotected (residues 1 −~38 and 102-140), although the N-terminal segment shows some heterogeneity. A continuous middle segment (residues ~39-101) is strongly protected by a systematically hydrogen bonded cross-β structure (60). The termini of the *α*S monomers are much more charged (net charge is −10 with 26 pieces of charged residues) than the middle (net charge is +3 with 13 pieces of charged residues). The number of H_2_O molecules bound to the protein (34–36) is 2.4(6)·10^2^ for the termini and it is 1. 5(3) · 10^2^ for the middle section, as summed for the individual amino acids. The same number for the whole *α*S is 4(1)·10^2^. Nominally, the terminal sequence binds 55% of the water molecules compared to the whole protein. Since the middle segment forms the β-sheet structure and therefore it is not involved so much in hydration, these hydration ratios can explain that the mobile hydration of the *α*S amyloid fibrils is approximately half of that of the *α*S monomers.

## Conclusions

*α*S variants are the studied proteins, namely WT *α*S and its A53T point mutant, in monomer, oligomer, and amyloid degrees of polymerization. Our earlier experimental results on aqueous solutions of these two *α*S variants (1) are reinterpreted. The bases of our experimental results are the temperature dependences of the number of mobile hydration ^1^H nuclei generating narrow-lines of two-component wide-line ^1^H NMR spectra (the MDs) and the thermodynamic interpretation of resulting melting diagrams (2,3,28). The two-component spectra originate from the different molecular mobility of water molecules bound to the protein also and not to other water molecules only. The experimentally determinable essential parameters of the thermodynamic or energy-based interpretation are directly measurable attributes, such as the start of motion of water molecules, start and endpoint of the constant-hydration sections in the melting diagrams (hydrophobic regions), the ratio of ordered or disordered regions and their measure. These are calculable also. The derivative of the melting diagram can be described analytically. It provides energy distributions of potential barriers hindering the motion of water molecules bound to protein molecules also in aqueous solutions of distilled water.

The above attributes of the two *α*S monomers are given first, and then similar characteristics of *α*S oligomer and *α*S amyloid variants of the same molecules are provided. The measurable molecular properties of these molecules could be quantified. The two *α*S monomers are nearly identical in their primary structure. These systems have not been investigated to such a deep level before. Comparisons can be made between the properties of *α*S monomers, oligomers, and amyloids. So, unlimited, promising possibilities are provided to explore the nature and causes of polymerization, or fibril forming processes also.

It could be demonstrated by direct experimental results that *α*S monomers are intrinsically disordered proteins with 33(4)% of their solvent accessible surface determined by secondary structures. That is, the monomers are more compact than it is expected for random coils. At the lowest potential barriers, the monomers have approximately monolayer coverage with mobile hydration water, and they are therefore functional. Their solvent-accessible surface is highly heterogenic regarding protein-water interactions. The mutant A53T *α*S monomer has a more open structure than the WT variant with more amino acid residues exposed to the solvent and therefore with a higher level of hydration than WT *α*S. The *α*S monomers have the highest ratio of solvent-exposed hydrophilic groups compared to the *α*S oligomer or *α*S amyloid forms. The comparison of the initial level of *α*S monomer hydration with amino acid hydration data available in the literature showed that every possible hydrogen bond is realized, confirming open structure again. This open nature involves that considering interaction energies, *α*S monomers interact with water most diversified.

One single potential barrier is responsible for the motion of the initial amount of mobile hydration water molecules in *α*S oligomers, which are therefore homogeneous in their bonding properties. *α*S oligomer forms have a uniform general surface; there is only one type of *α*S oligomers below the cubic melting-diagram section. The hydration of the *α*S oligomers is only slightly lower than that of the *α*S monomers, in the constant mobile hydration region. The wide, constant melting-diagram region is characteristic of globular structure (3), it is produced most possibly by β-sheets. 66(2)% of the *α*S oligomer variants’ solvent-accessible surface is assumed to be ordered, the *α*S oligomer states are mostly folded. The mobility of water in the *α*S oligomeric hydration shell is more limited than for the *α*S monomers or *α*S amyloids. Close to the melting point of bulk water, the melting diagrams of *α*S oligomers reflect their variety in size as a very heterogenic potential barrier distribution in a wide potential-barrier energy range, compared to melting diagrams of globular proteins.

The water molecules in the hydration shell of the *α*S amyloids can be mobilized the most easily compared to the case of *α*S monomers or *α*S oligomers. The hydration of the *α*S monomers is substantially higher in the constant mobile hydration region than in the case of the *α*S amyloids. The identical melting diagrams of the WT and A53T *α*S amyloid variants show that their structural features are independent of their sequence. The structure of the *α*S fibril exhibits many stabilizing features of the amyloid fold, including steric zippers involving the hydrophobic side chains and hydrophobic packing (66). Wide-line ^1^H NMR cannot see the water inside the solvent space within the *α*S fibrils as mobile, and this can be the cause of the low level of mobile hydration for *α*S amyloids. The potential barrier distribution affecting the motion of mobile hydration water is mainly heterogeneous in the *α*S amyloids, due to the abundant solvent-exposed side chains, the motion of which have bigger amplitudes than in the *α*S monomers (67). *α*S fibrils are disordered except for the core structure (67). In agreement with this fact, we found that the amyloid form of *α*S can be considered as disordered to 75(3)% regarding the nature of its solvent-accessible surface.

The initial amount of mobile water approximately equals in *α*S monomers and *α*S oligomers, while the *α*S amyloids have several times less such hydration moving at the lowest potential barriers. During the formation of *α*S oligomers or *α*S amyloids, approximately half of the mobile hydration water quantity, measured around *α*S monomers, is lost above the constant mobile hydration section, as the comparison of the melting diagrams of the three *α*S forms shows. These water molecules vanishing from the mobile hydration shell due to the aggregation of the monomers are connected to hydrophilic parts of the solvent accessible protein surface. The aggregation buries a significant part of the hydrophilic amino acid residues, and these become inaccessible to the solvent water. The buried amino acid residues form most possibly the structural fragments holding the *α*S oligomers and the *α*S amyloids together. These amino acid residues are presumably parts of the β-sheets.

## Author contributions

M.B. and K.T. conceived of the presented idea, devised and planned the experiments. M.B. carried out the experiments, processed the experimental data, and performed the analysis. M.B. and K.T. drafted the manuscript and designed the figures. P.T., Á.T. and K-H.H. helped supervise the project.

## Acknowledgements

Eszter Ságiné Házy is thanked for the preparation of the *α*-synucleins and for interpreting the initial results. The authors declare no conflicts of interest. This work was supported by the FWO Odysseus grant G.0029.12 (P.T.) from Research Foundation Flanders (FWO), the Korea-Hungarian Joint Laboratory program 2010-88343 (K-H.H., P.T.) from the Korea Research Council of Fundamental Science and Technology (KRCF), the National Research Council of Science and Technology (NST) grant NTM2231712 (K-H.H., P.T.), and grants K124670 (P.T.), K125340 (Á.T.).

